# Protein-Nucleic Acid Interactions for RNA Polymerase II Elongation Factors by Molecular Dynamics Simulations

**DOI:** 10.1101/2022.01.28.478254

**Authors:** Adan Gallardo, Brandon M. Bogart, Bercem Dutagaci

## Abstract

RNA polymerase II (Pol II) forms a complex with elongation factors to proceed the elongation stage of the transcription process. In this work, we studied elongation factor SPT5 and explored protein nucleic acid interactions for the isolated systems of KOW1 and KOW4 domains of SPT5 with DNA and RNA, respectively. We performed molecular dynamics (MD) simulations using three commonly used force fields that are CHARMM c36m, AMBER ff14sb and ff19sb. These simulations showed that most of the protein-nucleic acid interactions in the native state were retained with an increased electrostatic binding free energy for all force fields used. RNA was found highly dynamic with all force fields while DNA had relatively more stable conformations with the AMBER force fields compared to CHARMM. Furthermore, we performed MD simulations of the complete elongation complex using CHARMM c36m force field to compare the dynamics and interactions in the isolated systems. Similar strong KOW1 and DNA interactions were observed in the complete elongation complex simulations and DNA was further stabilized by a network of interactions involving SPT5-KOW1, SPT4 and rpb2 of Pol II. Overall, our study showed that the accuracy of force fields and the presence of the entire interaction network are important for elucidating the dynamics of protein-nucleic acid systems.

## INTRODUCTION

RNA polymerase II (RNA Pol II) synthesizes RNA in three main steps that are initiation, elongation, and termination^1^. Numerous biochemical^2-6^ and structural^7-11^ studies have added to understanding the mechanisms of these steps, demonstrating that other factors are involved in transcription. During initiation, general transcription factors are associated with RNA Pol II that forms the initiation complex^9, 11^. This complex dissociates as the RNA Pol II escapes the promoter site and forms the elongation complex with the basal elongation factors^7-8, 12^ to allow transcription with proper processivity. Structure of the transcription initiation complex has been well studied for the last decade^9, 11^ whereas the structure of elongation complex has only recently been revealed^7-8, 12^. The elucidated structure showed that elongation factors, SPT4 and SPT5, are bound to RNA Pol II to form DNA and RNA clamps at the exit sites suggesting that the interactions of the elongation factors with DNA and RNA are important for the elongation mechanism.

Protein-nucleic acid interactions are not only important for the elongation step of transcription but also in many fundamental biological processes like replication, translation and gene regulation. Thus, research on understanding protein-nucleic acid interactions became an emerging field. X-ray crystallography and cryo-EM provide structural information on protein-nucleic acid systems. In addition to the structural biology techniques, many other experimental techniques including Förster resonance energy transfer (FRET)^13-14^, nuclear magnetic resonance (NMR)^15-16^, mass spectrometry^17-18^, small angle X-ray scattering (SAXS)^19^, electrophoretic mobility shift assay (EMSA)^20-22^, footprinting^20-21^, crosslinking^21, 23^ and chromatin immunoprecipitation (ChIP)^20-21, 24-25^ focus on proteins-nucleic acids interactions and their dynamics during the biological processes. In addition to these studies, computational approaches based on molecular dynamics (MD) simulations have provided important insights on the dynamics of protein-nucleic acid interactions at the atomistic details^26-32^. MD simulations are used to obtain mechanistic understanding of the roles that protein-nucleic acid interactions play in biological processes^27-28, 32-33^. Studies have also shown the importance of force field choice in simulations to reproduce experimental values with high accuracy as they demonstrated that imperfections in the results can occur^29, 34-35^. The most widely used force fields are CHARMM^36-39^ and AMBER^40-43^ since they are largely validated based on various experimental data. Despite the extensive validation, there have been inaccuracies observed with these force fields in DNA^31, 44-45^ and RNA^34,46^ dynamics, diffusion calculations^29^ and protein-nucleic acid interactions^29, 47^. These results suggest that the force fields may generate different results on protein-nucleic acid systems and interpretation of the results should be done with care and consideration of possible simulation artifacts.

In this study, we performed MD simulations of elongation factors with nucleic acids to explore the differences among force fields in generating nucleic acid and protein dynamics and their interactions. We isolated the KOW1 and KOW4 domains of SPT5 and probed their interactions with DNA and RNA, respectively. We used three different force fields, which are CHARMM c36m, AMBER ff14sb and ff19sb. We observed that the interactions between protein and nucleic acids were retained in all force fields. However, force fields effected the flexibility of the DNA chains differently as the CHARMM force field displayed much higher fluctuation compared to both AMBER force fields. This suggests that AMBER force fields have larger stabilization effects on the intra-molecular interactions of DNA comparing to CHARMM. Furthermore, we used CHARMM c36m to simulate the whole elongation complex that includes RNA Pol II, SPT4, SPT5, upstream and downstream DNA, and nascent RNA. DNA fluctuations were observed in smaller amounts comparing to the isolated KOW1-DNA complex suggesting that DNA is stabilized not only by interactions with KOW1 but by an interaction network of Pol II and elongation factors in the whole complex.

## METHODS

### MD simulations of KOW1-DNA and KOW4-RNA systems

We used the elongation complex structure with PDB ID 5OIK^7^. For the KOW1-DNA systems we selected the SPT5 residues of 273-399 and DNA residues of 1-12 for the non-template and 32-43 for the template chains that are referred as DNAA and DNAB, respectively, in this paper. For the KOW4-RNA system, SPT5 residues of 536-646 and RNA residues of 31-38 were selected. We isolated those systems from the complete complex and solvated them in a cubic box with a 10 Å cutoff from each edge of the solvation boxes to prevent interactions with the periodic images. We added K^+^ ions to neutralize the systems. System preparation was performed by CHARMM-GUI server^48-50^ using CHARMM c36m^36, 39^, AMBER ff14sb^40^ and AMBER ff19sb^41^ for proteins. We used BSC1 parameters^42^ for DNA and OL3 parameters^43^ for RNA with the AMBER protein force fields. We used TIP3P parameters^51^ for the explicit water. We minimized the systems for 5000 steps with a tolerance of 100 kJ/mole. Systems were equilibrated for 625 ps with increasing temperature from 100 to 300 K with constraints at the backbone and side chains of proteins with force constant of 400 and 40 kJ/mol/nm^2^, respectively. We, then performed 1 μs of three replicates of MD simulations for each system with a total simulation time of 18 μs. KOW1-DNA system was simulated for four replicates since one of the replicates resulted in partial unfolding for the protein (see RMSD in Fig. S1) that we excluded from the rest of the analysis. Periodic boundary conditions were used with the particle mesh Ewald algorithm for the long-range electrostatic interactions. Lennard Jones interactions were switched between 1.0 to 1.2 Å for the simulations with CHARMM force field and 1.2 Å was used as a cutoff for the simulations with the AMBER force fields. Bonds with H atoms were constraint using the SHAKE algorithm. Simulations were run using Langevin thermostat at 303.15 K and with a friction coefficient of 1 ps^-1^ Time step was 1 fs for the equilibration and 2 fs for the production runs and trajectories were saved at every 20 ps. Simulations were performed using OpenMM^52^ on GPU machines.

### MD simulations of RNA Pol II elongation complex

The complete elongation complex with the PDB ID of 5OIK was used for the simulations. We modelled missing residues of the trigger loop (TL) using MODELLER program^53^. The system was solvated in a cubic box using a cutoff of 9 Å from each edge with a total box size of 176.5 Å. The system was prepared using MMTSB package^54^ in conjunction with the CHARMM software version 45b2^55^. The system was minimized and equilibrated as described above. We performed 200 ns of production run for each of the three replicates. Simulation details were the same as the small protein-nucleic acid systems described above. We used CHARMM 36m force field with TIP3P water parameters.

### Analysis of the simulations

The RMSD, RMSF, distance maps and contact analysis were performed using the MDAnalysis package^56^. For protein, C_α_-RMSD was calculated after superimposing the C_α_ and C_β_ atoms and for RNA and DNA, heavy atom RMSD was calculated after superimposing the heavy atoms. RMSF values were calculated for C_α_ atoms of proteins and P atoms for nucleic acids after superimposing the C_α_-C_β_ atoms and heavy atoms to the average structures, respectively. Distance maps were generated for the initial experimental structure and for the simulations averaged over the trajectories and plotted for distances lower than 10 Å. Contacts between protein and DNA were averaged over the trajectories for each residue pairs that were within 5 Å distance. We calculated the free energy profiles for protein-nucleic acid distances of the simulation trajectories using weighted histogram analysis method^57-58^. The RMSD analysis was performed for the whole trajectories while all the other analyses were performed for the last 800 ns of the simulations for the KOW1-DNA and KOW4-RNA systems, since the fluctuations of C_α_-RMSD decreased after the first 200 ns simulation (Fig S1-2). The analysis for the complete RNA Pol II simulations were performed for the last 150 ns since the RMSD of the KOW1 and KOW4 domains became stable after 50 ns of the simulations (Fig S3).

We performed energy calculations using Molecular Mechanics Generalized Born and Surface Area (MMGB/SA) approach^59-60^ using the MMTSB package^54^. We calculated total, electrostatic and van der Waals (vdW) binding energies for protein and DNA/RNA pairs without including the entropic contribution. For the energy calculations of the isolated KOW1-DNA and KOW4-RNA systems, we obtained 1,000 conformations extracted at every 2.4 ns from three replicates of simulations for each force fields and applied 500 steps of minimization for each conformation before the energy calculation. For the complete elongation complex, we employed energy calculations onto the total of 900 frames extracted at every 0.5 ns. For the native state, we extracted the KOW1-DNA and KOW4-RNA from the original structure and employed energy calculation after adding hydrogen atoms and applying 5000 steps of minimization. All the minimizations were performed with Adopted Basis Newton-Raphson algorithm. We calculated energies using CHARMM c36 force field.

Hydrogen bond analysis were performed for the Watson-Crick base pairs such that the hydrogen bonds were counted between N1 of adenine and N3 of thymine, N6 of adenine and O4 for thymine, O6 for guanine and N4 of cytosine, N1 of guanine and N3 of cytosine, N2 of guanine and O2 of cytosine. The average number of hydrogen bond counts for the DNA pair was calculated over the trajectories of three replicates. Hydrogen bond analysis was performed using MDAnalysis package^56^.

## RESULTS

In this study, we explored the protein-nucleic acid interactions and dynamics for the KOW1 and KOW4 domains of SPT5 elongation factor using AMBER and CHARMM force fields. We isolated the KOW domains and the interacting nucleic acids for the simulations as seen in Fig. 1. We analyzed the simulations in terms of direct interactions between proteins and nucleic acids and the dynamics of both components. We focused on the differences between the simulations with varying force fields to understand the strengths and weaknesses of the force fields for the isolated systems. We, then simulated the complete complex with the CHARMM force field and compared the results with the isolated systems. The results are presented in the following sections.

**Figure 1.**
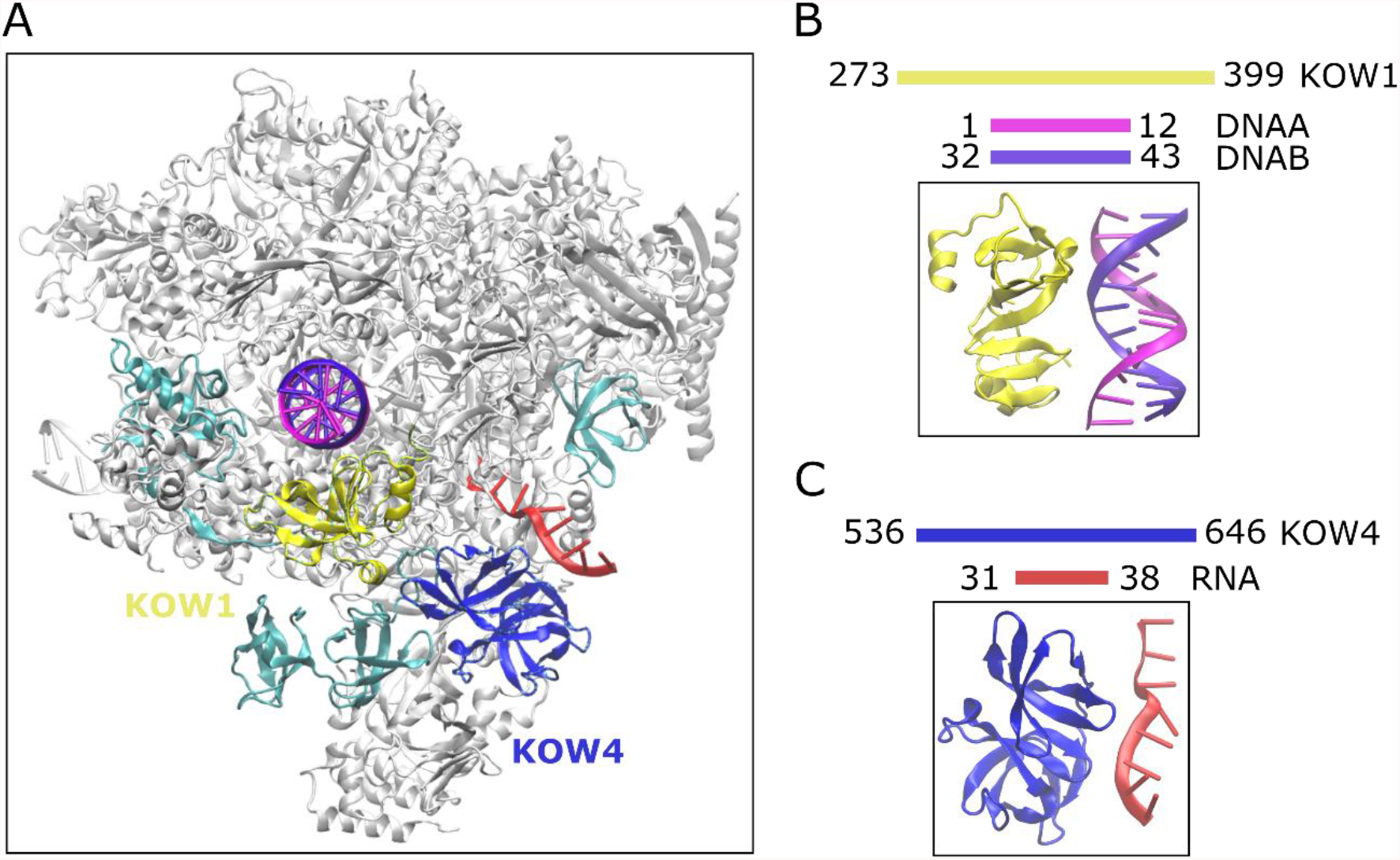
The structures and selections of the simulated systems. (A) the complete elongation complex with the following color codes: KOW1 yellow, KOW4 blue, RNA red, DNAA (non-template) magenta, DNAB (template) violet, other SPT5 domains cyan, and the rest of the complex gray. Selection of the residues and the structures for (B) KOW1-DNA and (C) KOW4-RNA are shown. The figure was generated using the structure with the PDB ID of 5OIK.

### KOW1-DNA interactions

The simulations of KOW1 and DNA were performed with three force fields. CHARMM c36m provided a large fluctuation in RMSD values for DNA chains comparing to the simulations with AMBER force fields (Fig. S1). Fig. 2A shows the RMSF profiles for protein (KOW1) and DNA, in which DNA chains show larger RMSF values with CHARMM c36m. For the protein, the RMSD and RMSF profiles are similar for the CHARMM and AMBER force fields although there are larger fluctuations for the residues from 340 to 365 for the AMBER simulations. This is due to the partial unfolding of the beta sheet region in AMBER simulations, while these beta sheets are retained to some extend in CHARMM c36m simulations.

**Figure 2.**
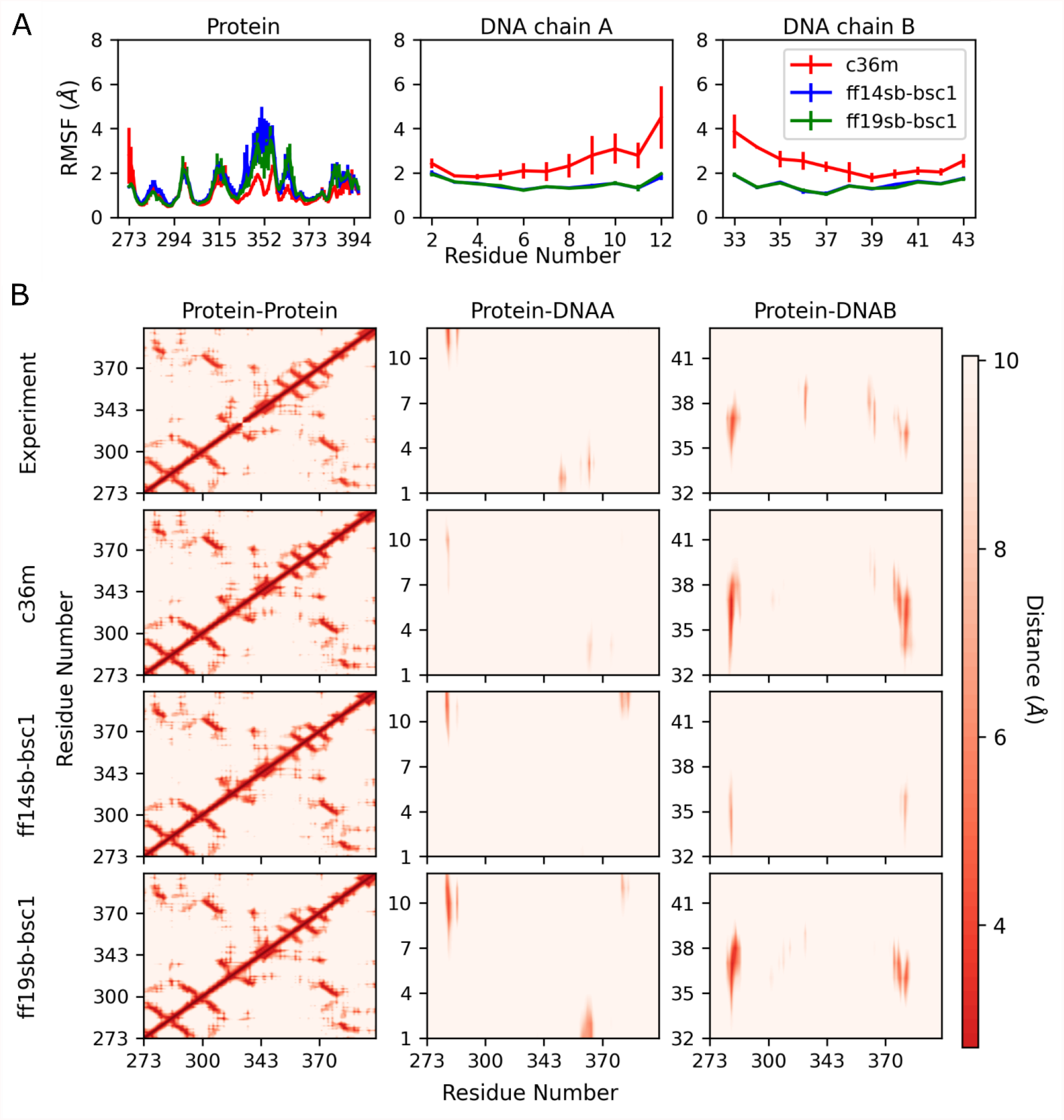
The RMSF and distance maps analysis for the simulations of the KOW1-DNA system with CHARMM c36m, AMBER ff14sb and AMBER ff19sb force fields. (A) RMSF values were calculated for protein, DNAA, and DNAB, (B) distance maps between protein-protein, protein-DNAA and protein-DNAB residues were calculated from the distances for the experimental structure (PDB ID:5oik) and average distances over the trajectories.

In Fig. 2B, we compared the distance maps of native structures for KOW1-DNA systems with the distances averaged over the simulations. Protein-protein interactions are mostly retained during the simulations with CHARMM and AMBER ff19sb force fields, while some of the interactions for the loop regions are lost in the simulations with AMBERff14sb. The two loops with residues, 335-340 and 285-289 have close distances at the native structures, while AMBER ff14sb simulations produced larger distances for these loops. Protein-DNA distances showed larger changes between force fields than the protein-protein distances in the simulations. Protein-DNAA interactions were partly retained with three force fields. DNAB shows strong interactions with protein for CHARMM and AMBERff19sb, although some distances were larger especially for the residues K317 and R365. However, most of the distances were increased with AMBERff14sb. This is partly due to one replicate of the AMBER ff14sb simulations, in which DNA and protein interactions were mostly lost comparing to the native state (Fig S4) and this affected the average distances over the replicates. Overall CHARMM and AMBER ff19sb simulations provided more consistent average distance maps with the native structures, while AMBER ff14sb showed increased distances for protein-protein and protein-DNA distance maps.

To have a closer look on how protein and DNA are interacting, we plotted the free energy landscapes for the selected distances of R283 and R365 with the DNAB residues 37 and 38, respectively. We selected R283 and R365 for this analysis since they have close distances with the DNA residues in the initial structure (Fig. 2B). Fig. 3 shows the free energy plots for the selected distances and the most probable structures. In one of the AMBER ff14sb simulations, protein and DNA lost most of the interactions (see Fig S4) and, therefore, we excluded this trajectory in the analysis shown in Fig 3. CHARMM c36m simulations provided an energy landscape with multiple minimum energy conformations while AMBER force fields provided only one minimum energy conformations. DNA chains with AMBER force fields were more stable as seen in the RMSF curve and Fig. 3 also shows that the conformations from AMBER simulations retained most of the base-paired interactions. On the other hand, in CHARMM simulations based-pair interactions are lost to some extent. Distance maps between DNA chains in Fig S5 also show that DNAA-DNAB distances at the end of the chains were increased in CHARMM c36m simulations suggesting that base-pair interactions for terminal residues are less stabilized comparing to the AMBER force fields. To quantify the base-pair interactions, we calculated the average number of hydrogen bonds for Watson-Crick pairs and found that CHARMM c36m simulations provided only around the half of the hydrogen bond count obtained by AMBER simulations (Table 1).

**Table 1.** Average number of hydrogen bonds.

|  | <sup>1</sup>Number of H-bonds |
| --- | --- |
| <sup>2</sup>Experiment | 24 |
| C36m | 9.78±0.02 |
| ff14sb-bsc1 | 19.13±0.02 |
| ff19sb-bsc1 | 19.25±0.02 |
| C36m<br>(Pol II-EC complex) | 14.85±0.02 |

<sup>1</sup>Number of H-bonds between Watson-Crick pairs was calculated for each frame and then averaged over the trajectories. <sup>2</sup>Number of H-bonds was calculated from the initial structure that was extracted from the structure with PDB ID:50IK.

**Figure 3.**
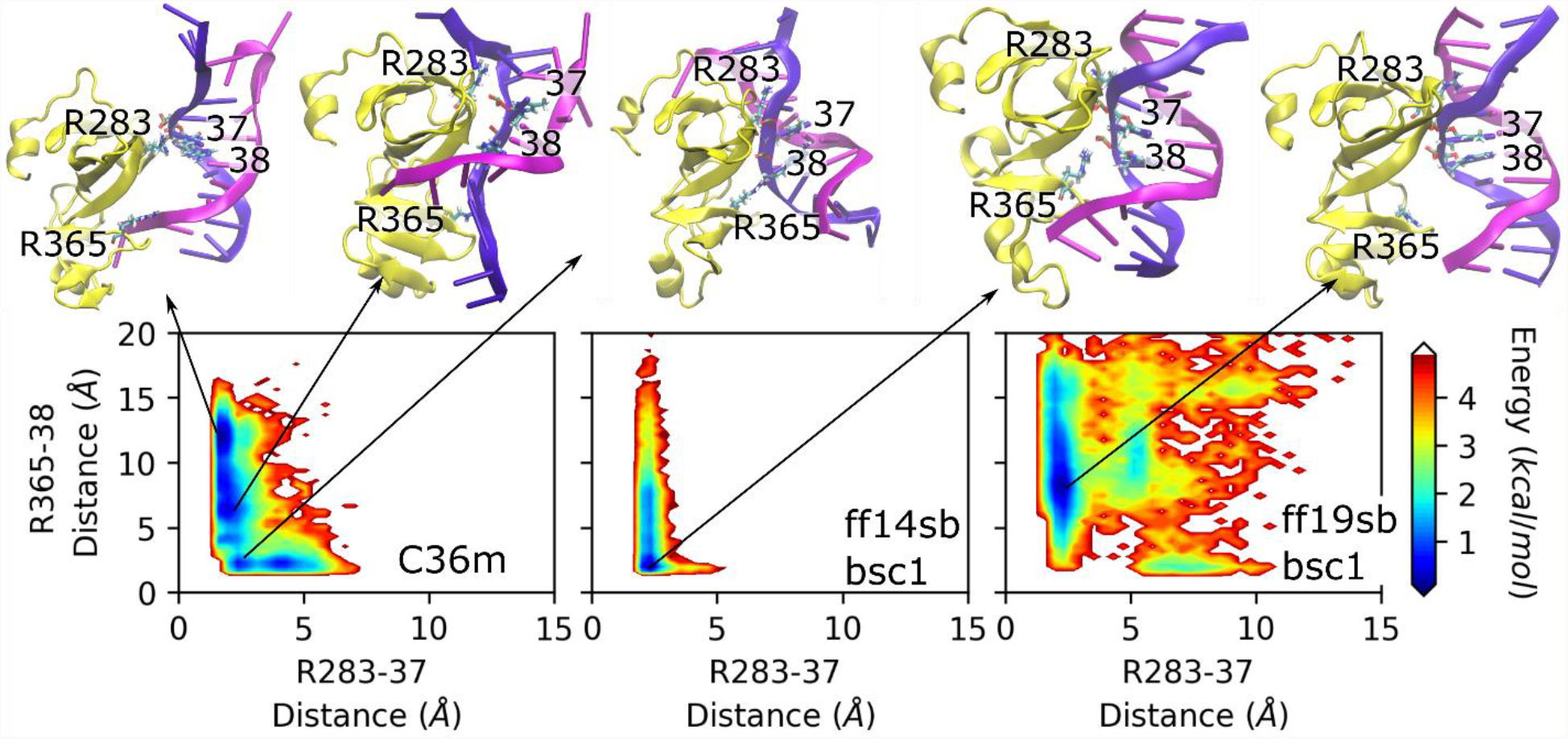
Free energy landscapes for the distances between R283 of protein and residue 37 of DNAB (x-axis) and R365 of protein and residue 38 of DNAB (y-axis). On the top, the snapshots at the minimum energy conformations are shown. Proteins were superimposed to the native state.

To understand electrostatics and vdW energy contributions to the DNA and protein binding energies, we performed MMBG/SA calculations (see details in Methods). All three force fields provided overestimation of both electrostatics and vdW binding energies while in CHARMM c36m simulations, overestimation was far greater compared to the AMBER force fields (Table 2). Overall, AMBER force fields were stabilizing both DNA base-pair and protein-DNA interactions; CHARMM c36m force field stabilized the protein-DNA interactions to a larger extent while it resulted in more fluctuations in DNA chains with decreased base pair interactions.

**Table 2.** Binding free energies calculated using MMGB/SA method.

| KOW1-DNA |  |  |  |
| --- | --- | --- | --- |
|  | Total binding energy | Electrostatic binding energy | vdW binding energy |
| <sup>1</sup>Experiment | -21.56 | -1364.92 | -14.68 |
| C36m | -103.86 ± 0.76 | -2134.27 ± 5.25 | -60.11 ± 0.48 |
| ff14sb-bsc1 | -71.86 ± 0.76 | -1765.30 ± 10.36 | -45.96 ± 0.46 |
| ff19sb-bsc1 | -79.76 ± 0.45 | -2031.15 ± 5.93 | -46.29 ± 0.32 |
| C36m<br>(Pol II-EC<br>complex) | -55.03 ± 0.52 | -1832.52 ± 6.28 | -34.36 ± 0.55 |
| KOW4-RNA |  |  |  |
|  | Total binding energy | Electrostatic binding energy | vdW binding energy |
| Experiment | -19.34 | -844.44 | -11.15 |
| C36m | -69.81 ± 0.56 | -929.34 ± 2.66 | -47.13 ± 0.37 |
| ff14sb-ol3 | -96.31 ± 0.69 | -1128.82 ± 3.96 | -75.34 ± 0.67 |
| ff19sb-ol3 | -101.67 ± 0.60 | -1114.53 ± 3.16 | -74.80 ± 0.36 |
| C36m<br>(Pol II-EC<br>complex) | -64.06 ± 0.57 | -985.71 ± 3.14 | -43.01 ± 0.35 |

<sup>1</sup>Energies were calculated for the initial structure that was extracted from the structure with PDB ID:50IK after minimization.

### KOW4-RNA interactions

KOW4 subunit of SPT5 protein has mostly well-defined secondary structure and smaller loop regions comparing to KOW1. This is retained for the simulations with all three force fields as RMSD values converged to less than 2 Å (Fig. S2) and RMSF plots (Fig. 4A) show only small fluctuations. However, for RNA, RMSD values were large and fluctuating for all the force fields and RMSF values were higher than proteins with larger error bars (Fig. 4A). Large fluctuations in RNA structure were expected as it does not have well-defined secondary structure.

**Figure 4.**
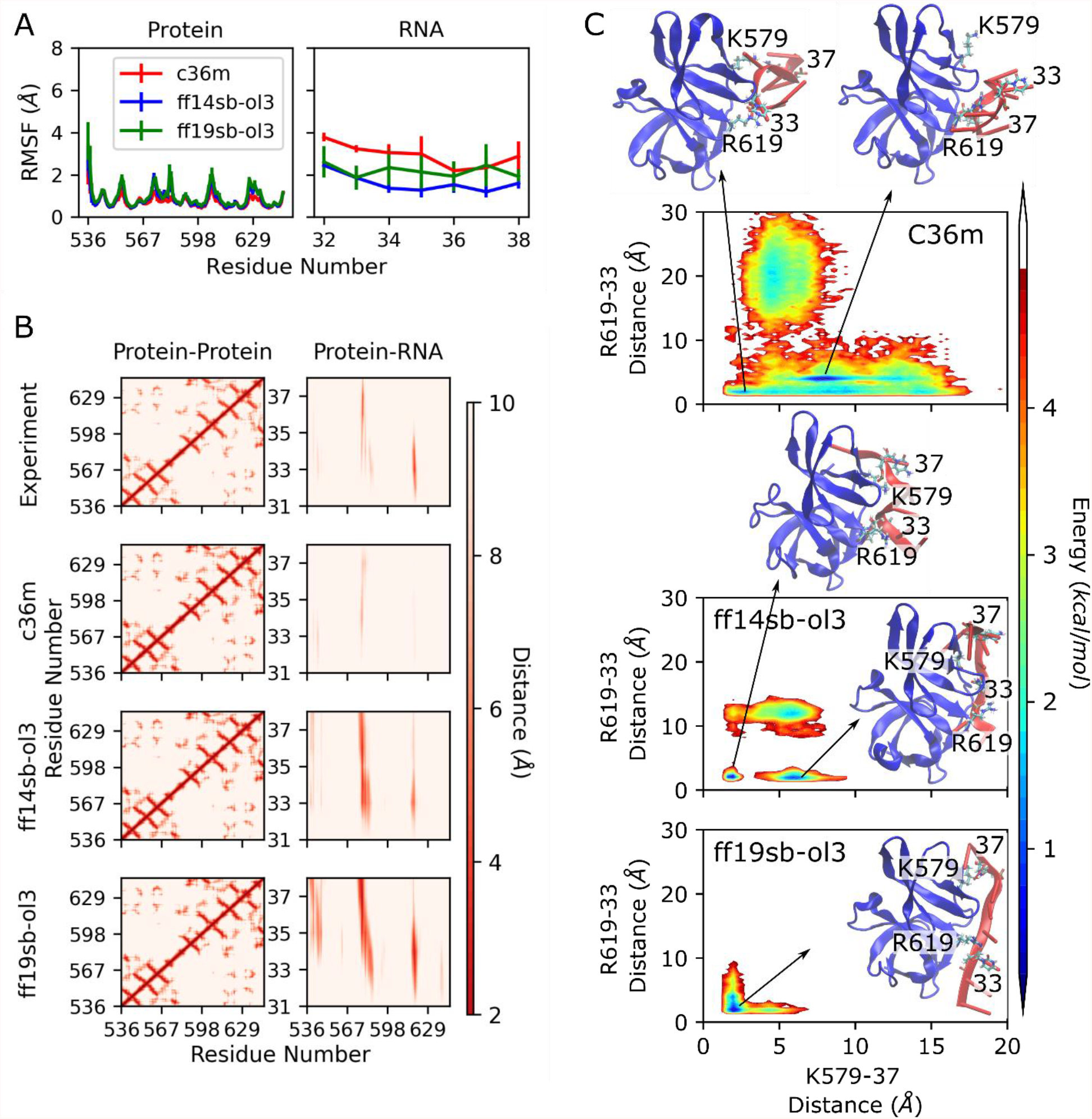
RMSF, distance maps, and free energy landscape analysis for the simulations of KOW4-RNA system with CHARMM c36m, AMBER ff14sb and AMBER ff19sb force fields. (A) RMSF values for protein and RNA were calculated, (B) distance maps calculated from the average distances over the trajectories between protein-protein and protein-RNA, (C) free energy landscapes for the distances between K579 of protein and residue 37 of RNA (x-axis) and R619 of protein and residue 33 of RNA (y-axis) along with the snapshots at the minimum energy conformations are shown.

Distance maps for the protein (KOW4) were also maintained in large extent during the simulations as protein structures were mostly preserved (Fig. 4B). However, the distance maps between protein and RNA change to some extent with force fields. RNA is in close contact with the residues of K579 and R619 of the protein in the experimental structure. Those interaction sites were also observed in all three force fields with some differences. AMBER force fields showed strong interactions of RNA with protein at the K579 and R619 residues, whereas CHARMM force field simulations provided longer average distances for the K579 and R619 residues. Fig. 4C shows the free energy plots for the distances of K579 and R619 with RNA residues of 37 and 33, respectively. Simulations with CHARMM force field showed a broader landscape with multiple minimum energies whereas simulations with AMBER force fields provided relatively narrower landscapes especially with ff19sb. Overall, RNA is more flexible in CHARMM simulations and its interaction sites with protein change frequently, while AMBER simulations mostly maintained the interaction sites.

Table 2 shows the binding energies for RNA and protein. Like KOW1-DNA interactions, KOW4-RNA interactions were also over-stabilized by the three force fields. In this case, AMBER force fields stabilized the protein-RNA interactions more than CHARMM force field, which is consistent with the distance maps that protein-RNA distances are smaller with AMBER force fields that imply stronger interactions.

Overall, the force fields were able to provide consistent structures for both KOW1 and KOW4. Protein-nucleic acid interactions were more difficult to reproduce by the force-fields, due to high flexibility in the DNA and RNA chains and differences in stabilization of base-pairs and electrostatic interactions in AMBER and CHARMM force fields.

### KOW1-DNA and KOW4-RNA interactions in the complete elongation complex

KOW1 and KOW4 are domains in SPT5 elongation factor that is associated with RNA Pol II during the elongation step. We performed simulations of the complete elongation complex to observe the changes in interactions of KOW1-DNA and KOW4-RNA when they are in complex with RNA Pol II. Fig. S3 showed that DNA chains were converged to lower RMSD values comparing to the isolated system (Fig. S1) using CHARMM force field, which indicates that the DNA chains were stabilized further in the complete complex comparing to the isolated system and gave closer conformations to the experimental structure. Fig. S3 shows that RMSD values for the RNA chain are also lower compared to the isolated KOW4-RNA complex (Fig. S2) while they are much higher than the DNA chains. Similarly, RMSF values were reduced in complete complex for both DNA and RNA chains (Figs 5A and 5C). For the proteins, KOW1 and KOW4, the RMSD and RMSF values are similar for the isolated and full elongation complex systems.

**Figure 5.**
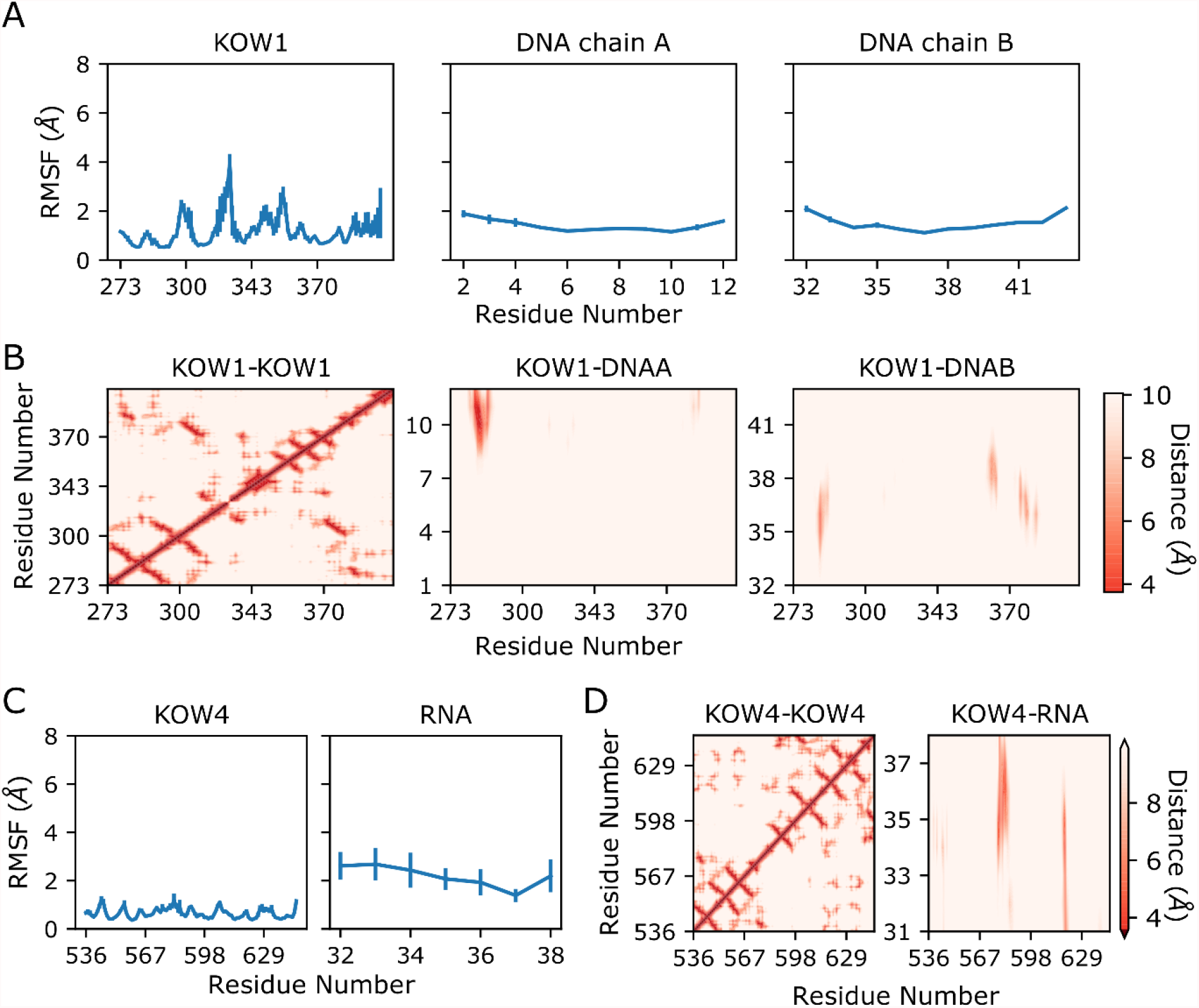
RMSF and distance maps for KOW1-DNA and KOW4-RNA from the simulation of the complete elongation complex with CHARMM c36m force field. (A) RMSF for KOW1 and DNA chains, (B) distance maps for KOW1-KOW1, KOW1-DNAA and KOW1-DNAB, (C) RMSF for KOW4 and RNA chains, (D) distance maps for KOW4-KOW4, KOW4-RNA.

The distance maps (Fig. 5B) showed that DNA and KOW1 were interacting strongly throughout the trajectories and interactions sites were mostly the same sites with the experimental structure (Fig. 2B). KOW4 and RNA distance map (Fig. 5D) was also similar to the map of the experimental structure (Fig. 4B). Table 2 shows that the binding energies for KOW1-DNA are closer to the experimental structure and less over-estimated compared to the isolated system, while the energies are similar to the isolated system for KOW4-RNA.

Fig. 6 shows the free energy landscape for the selected protein-DNA (Fig. 6A) and protein-RNA distances (Fig. 6B). Protein-DNA distances show a broad distribution like in the isolated CHARMM c36m simulations, however, there is one minimum energy conformation that is mostly populated. The minimum energy conformation shows that DNA chain retained base-pair interactions in larger extent compared to the isolated system. In agreement with this, hydrogen bond count between DNAA and DNAB chains were larger in the complete system (Table 1). Furthermore, we analyzed the number of contacts between the DNA chains and all the proteins in the elongation complex to observe if DNA has interactions with any other protein domain in addition to KOW1 that could contribute to the stabilization of the DNA chains. Table 3 shows that 45.63 % of the contacts of the DNA chains (the selected region in Fig 1) were observed with SPT5-KOW1 domain, while SPT4 domain and Pol II rpb2 subunit also have large number of contacts with DNA. These additional interactions may also contribute to the DNA dynamics such that the DNA chains are fluctuating less and have more stabilized base pair interactions. This suggests that simulating the isolated KOW1-DNA system may not be sufficient to cover the DNA dynamics that is affected by a network of interactions within the elongation complex as seen in Fig. 7.

**Table 3.**
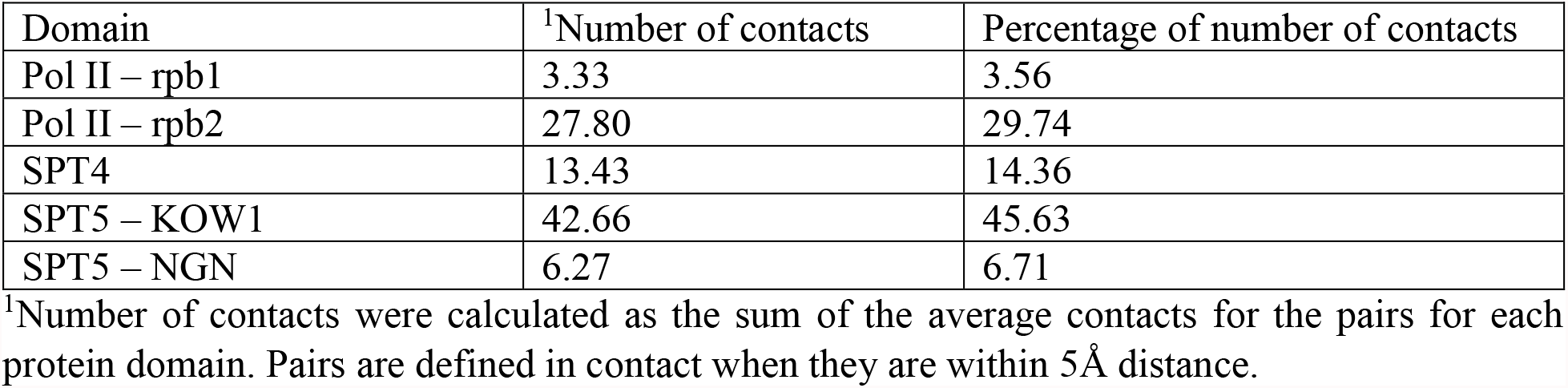
Number of contacts between DNA and protein in the elongation complex.

| Domain | <sup>1</sup>Number of contacts | Percentage of number of contacts |
| --- | --- | --- |
| Pol II – rpb1 | 3.33 | 3.56 |
| Pol II – rpb2 | 27.80 | 29.74 |
| SPT4 | 13.43 | 14.36 |
| SPT5 – KOW1 | 42.66 | 45.63 |
| SPT5 – NGN | 6.27 | 6.71 |

<sup>1</sup>Number of contacts were calculated as the sum of the average contacts for the pairs for each protein domain. Pairs are defined in contact when they are within 5Å distance.

**Figure 6.**
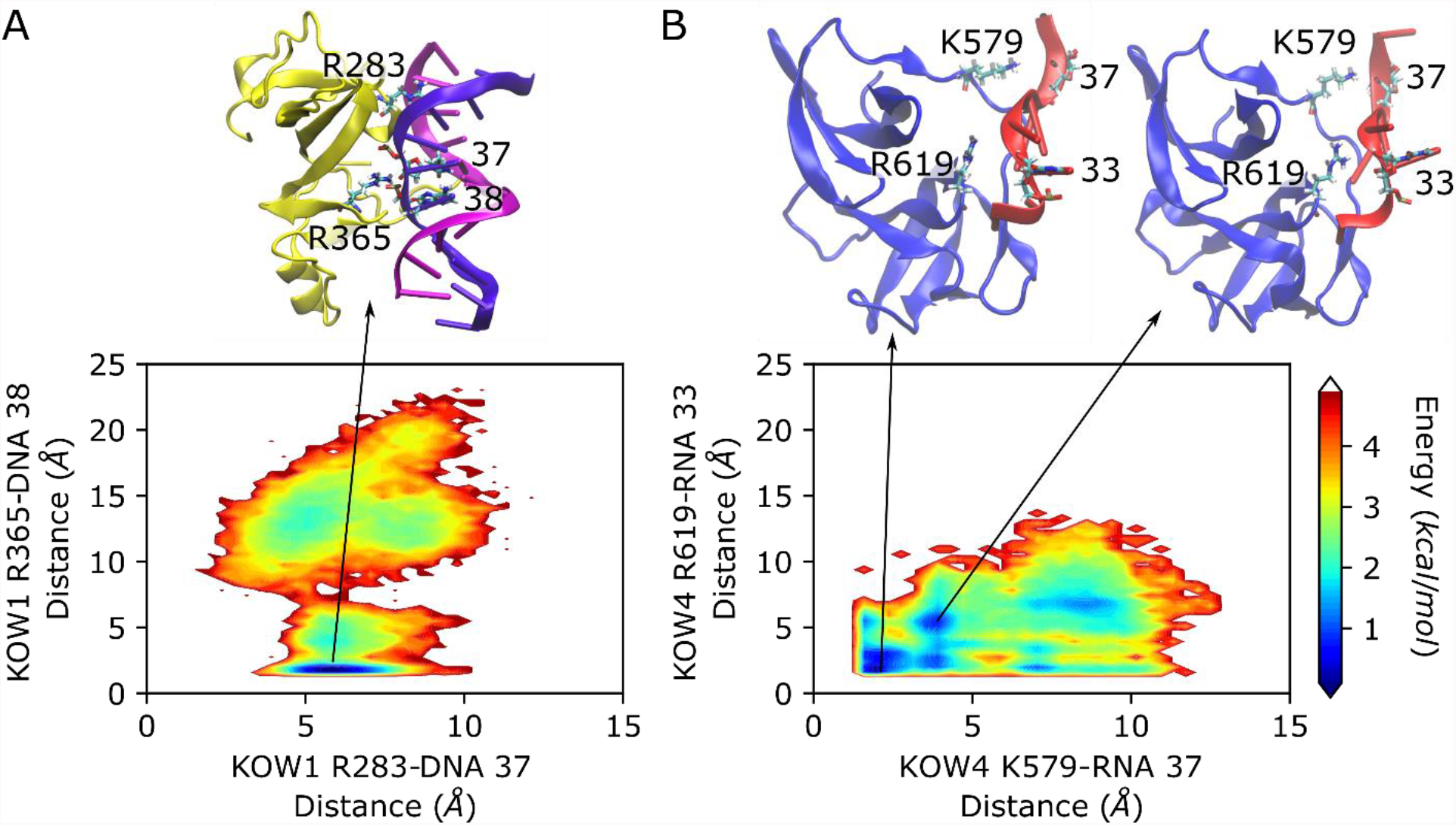
Free energy landscapes for the distances between KOW1 and DNA, and KOW4 and RNA. (A) R283 of KOW1 and residue 37 of DNAB (x-axis) and R365 of KOW1 and residue 38 of DNAB (y-axis); (B) K579 of KOW4 and residue 37 of RNA (x-axis) and R619 of KOW4 and residue 33 of RNA (y-axis); on the top are snapshots at the minimum energy conformations.

**Figure 7.**
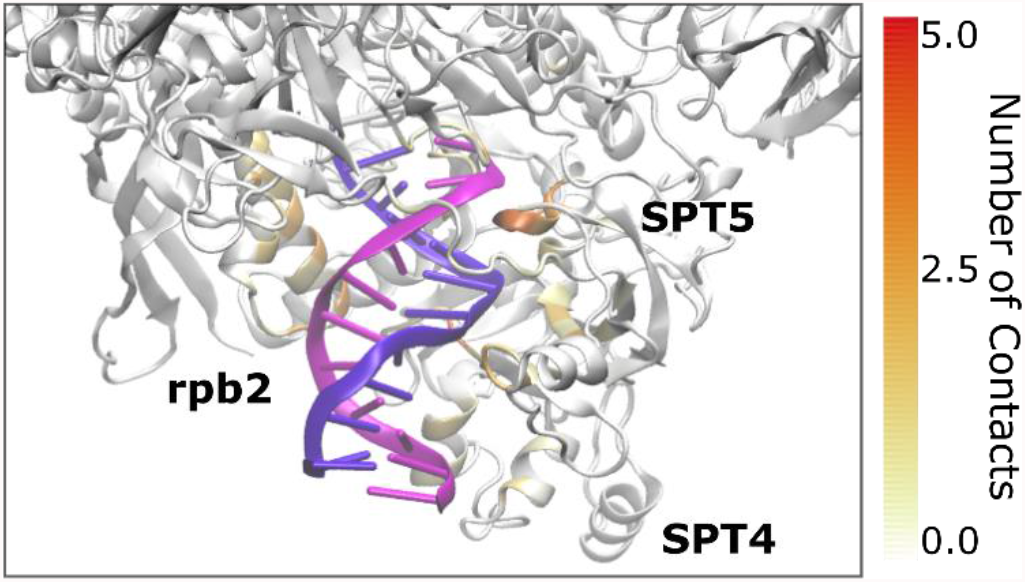
Number of contacts of DNA with the surrounding protein residues colored as depicted in the color bar. The elongation complex structure obtained at the minimum energy state in Fig. 6A was used.

Overall, the interactions of KOW1 and KOW4 with the nucleic acids are strong and stabilized in the full elongation complex similar to the isolated system. On the other hand, DNA fluctuations were reduced in the complete complex with an increased number of hydrogen bonds between the DNA pairs compared to the isolated system. This suggests that simulating DNA-protein systems in isolation may not cover the complete dynamic picture of the nucleic acids and the other interacting proteins should be included in the simulations to obtain higher accuracy in DNA dynamics.

## DISCUSSION

Protein-nucleic acid interactions are important for many biological processes and MD simulations are a powerful method to probe these interactions at the atomistic detail in μs time scales. One important part of this approach is the capability of the force field to maintain accurate results in the protein-nucleic acid interactions and in the dynamics of both components. CHARMM and AMBER are the two most used force fields for protein-nucleic acid systems. Although they went through several validations for the proteins^36, 39-41^ and nucleic acids^37-38, 42-43^, there are still imperfections reported for these force fields for protein-nucleic acid systems^29, 31, 47^. In our work, we used three recent force fields, AMBER ff14sb and ff19sb with bsc1 for DNA and ol3 for RNA and CHARMM c36m, to investigate the differences between force fields in production of the protein and DNA interactions. We observed that the electrostatic interactions between protein and DNA or RNA are over-estimated with both AMBER and CHARMM force fields. The overestimation of the charge-charge interactions was reported by previous studies as well for both standard AMBER and CHARMM force fields^28-29, 35, 47^. One remedy for accounting for the unrealistic electrostatic interactions was proposed to modify Lennard Jones parameters to balance pairwise interactions for the charged residues^29, 35, 47^. Alternatively, there are studies focused on water models, proposing modified water-solute interactions, which could reduce the overestimation of solute-solute interactions^61-63^.

Another result of our simulations is the difference in the stability of base pair interactions in AMBER and CHARMM force fields, with AMBER having larger stabilization effect on these interactions. For both AMBER and CHARMM force fields, we observed a smaller number of Watson-Crick hydrogen bonds than in the native state. This is consistent with what was found in the literature, that both standard AMBER and CHARMM force fields overestimate the stacking interactions that comes with the underestimation of the base pair interactions^44-45, 64^, while we found the underestimation is larger in CHARMM c36m simulations. Recent studies on improvement of RNA force fields suggest multiple modifications on vdW parameters and charges on nucleobase to balance stacking interactions with the base pairing strength^34, 46^ although transferability of these modifications into the DNA force fields was not studied. In addition, four sited water models that are better balancing between solute-solute and solute-water interactions were also known to provide improvement in the stacking energies^34, 64-65^.

The discrepancies in the protein nucleic acid interactions we observed in the isolated systems were reduced when we simulated the complete elongation system. The binding energies between the protein and DNA were still overestimated but closer to the experimental system compared to the isolated KOW1-DNA system. The base pair interactions between DNA chains were also improved in the complete elongation complex, but they are still underestimated comparing to the native state. Overall, the standard CHARMM c36m force field provides more accurate results as the system become more realistic and biologically relevant as a complex, but there are still imperfections in the protein-nucleic acid interactions produced by CHARMM c36m force field.

Our study altogether suggests that the protein nucleic acid interactions are captured well by both standard AMBER and CHARMM force fields, with some imperfections in the electrostatic interactions and base pair interactions that require attention for obtaining realistic simulations for protein-nucleic acid systems. A future direction could be to apply suggested modifications that includes scaling the water-protein interactions as well as modifying Lennard Jones interactions between the protein and nucleic acids to improve the accuracies of the force fields.

## CONCLUSION

Proteins and nucleic acids strongly interacted with each other during the simulations of KOW1 and KOW4 domains of SPT5 elongation factor with DNA and RNA using CHARMM and AMBER force fields. However, the interactions between protein and nucleic acids were over stabilized with both CHARMM and AMBER force fields compared to the native interactions. Simulations of the complete elongation complex, on the other hand, provided a more realistic environment and reduced the discrepancies observed in the isolated system to some extent. In addition, we observed that the DNA was stabilized by a network of interactions with the proteins in the Pol II elongation complex suggesting that isolated systems may not provide accurate dynamics for DNA. Future studies will focus on simulating the complete elongation complex rather than using the isolated systems and applying the modifications suggested in the literature into the standard force fields to explore the interactions of elongation factors with nucleic acids as well as the changes in DNA and RNA dynamics.

## AUTHOR INFORMATION

**Note**

The authors declare no competing financial interest.

## ACKNOWLEDGMENTS

We used the computational resources at the MERCED cluster at University of California Merced and at the National Science Foundation’s Extreme Science and Engineering Discovery Environment (XSEDE) facilities under the grant TG-BIO210145.

## SUPPORTING INFORMATION

Figures S1-5 are provided as Supporting Material.

## Notes

### Competing Interest Statement

The authors have declared no competing interest.

