## Supplementary Information for "Protein-Nucleic Acid Interactions for RNA Polymerase II Elongation Factors by Molecular Dynamics Simulations"

**This PDF file includes:**

Supplementary Figures S1 to S5

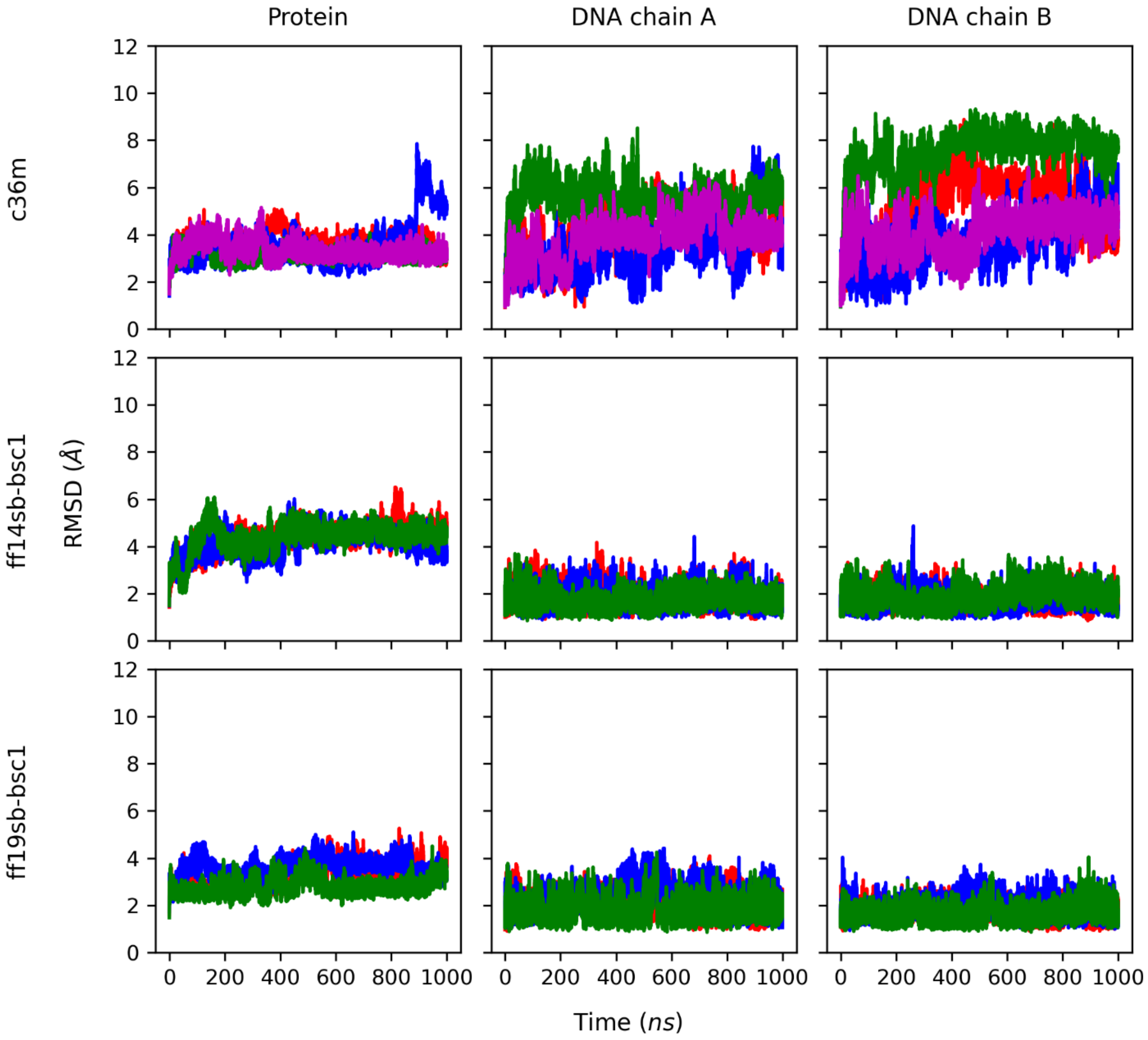


**Figure S1.** RMSD values for protein and DNA chains in KOW1-DNA simulations with CHARMM c36m, AMBER ff14sb-bsc1 and ff19sb-bsc1 force fields. RMSD values were reported for $C_\alpha$ for the protein and heavy atoms for the DNA chains. Different colors show the values for the three replicates for all the AMBER simulations and four replicates for the simulations with c36m.

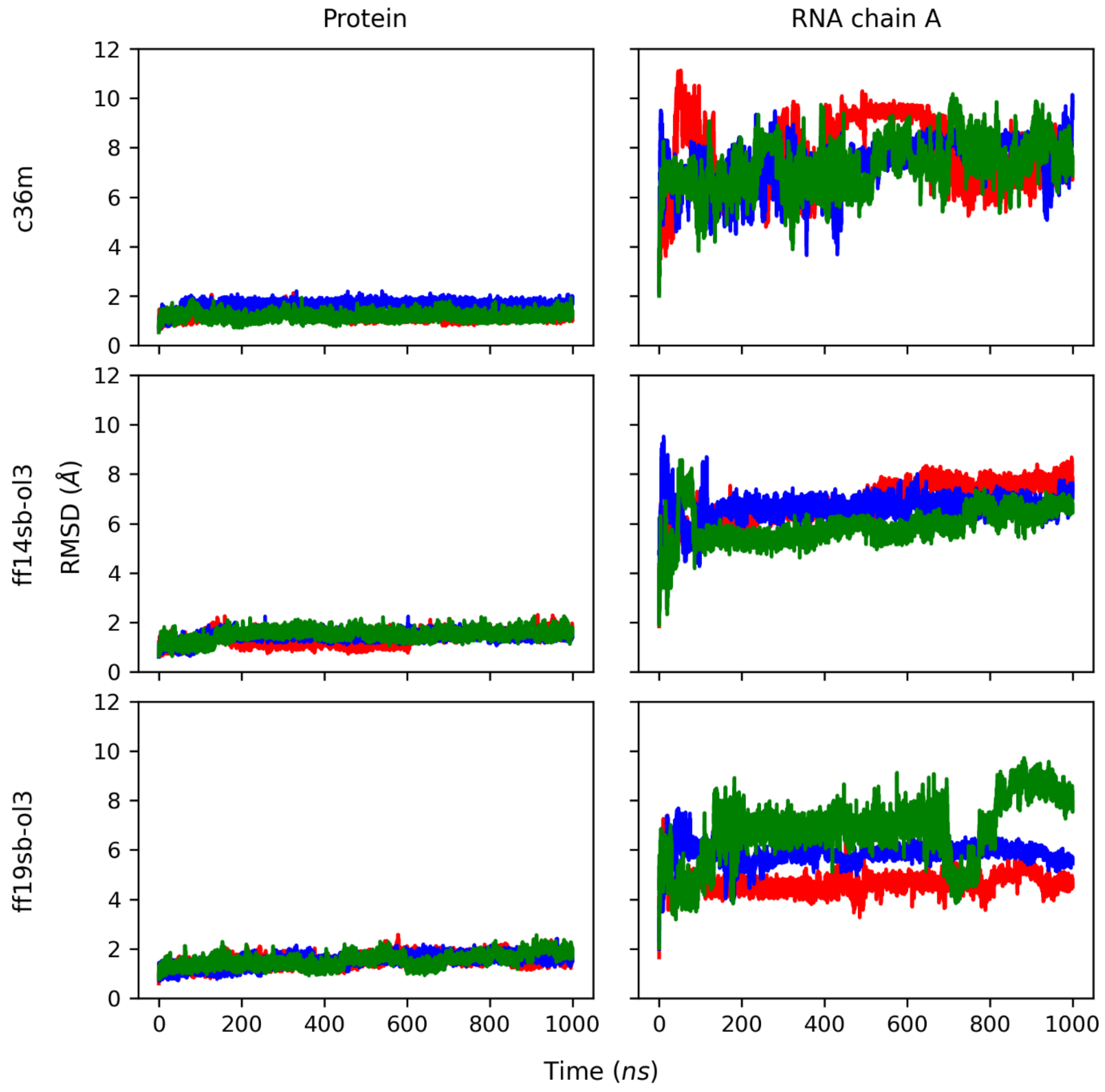


**Figure S2.** RMSD values for protein and RNA in KOW4-RNA simulations with CHARMM c36m, AMBER ff14sb-bsc1 and ff19sb-bsc1 force fields. RMSD values were reported for $C_\alpha$ for the protein and heavy atoms for the RNA. Different colors show the values for the three replicates.

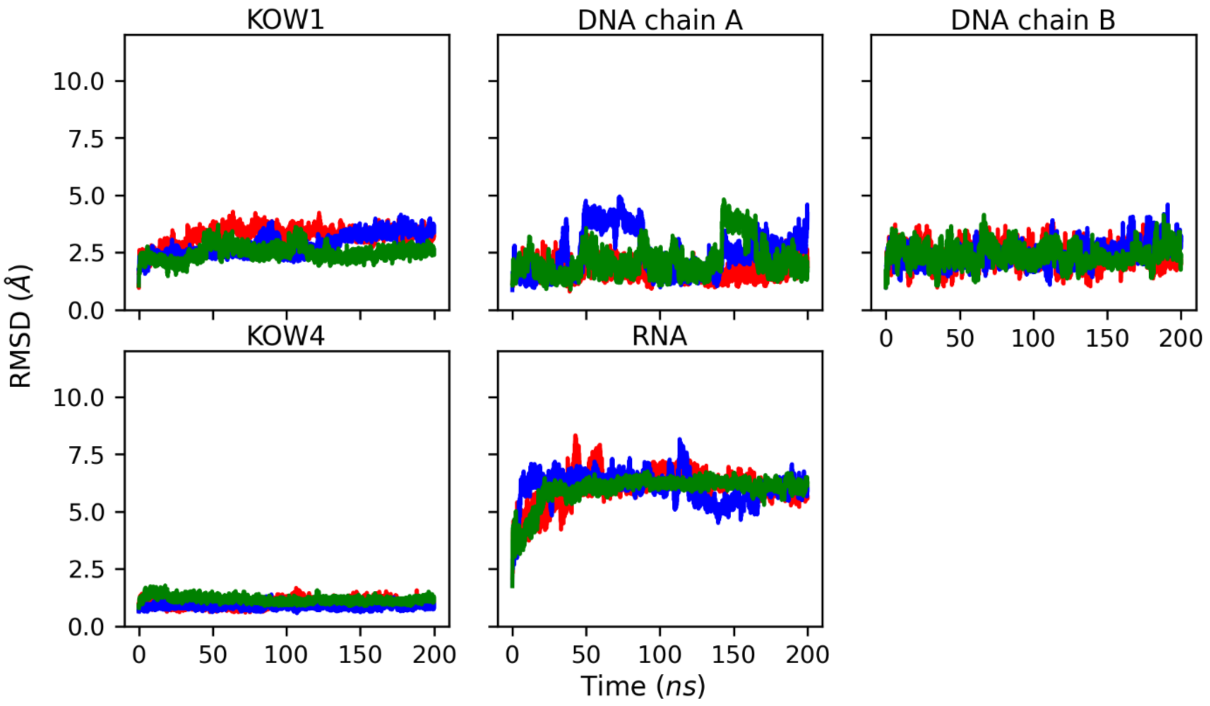


**Figure S3.** RMSD values for KOW1, KOW4, DNA and RNA chains in the complete elongation complex simulations with CHARMM c36m force field. RMSD values were reported for $C_\alpha$ for the protein and heavy atoms for the DNA and RNA chains. Different colors show the values for the three replicates.

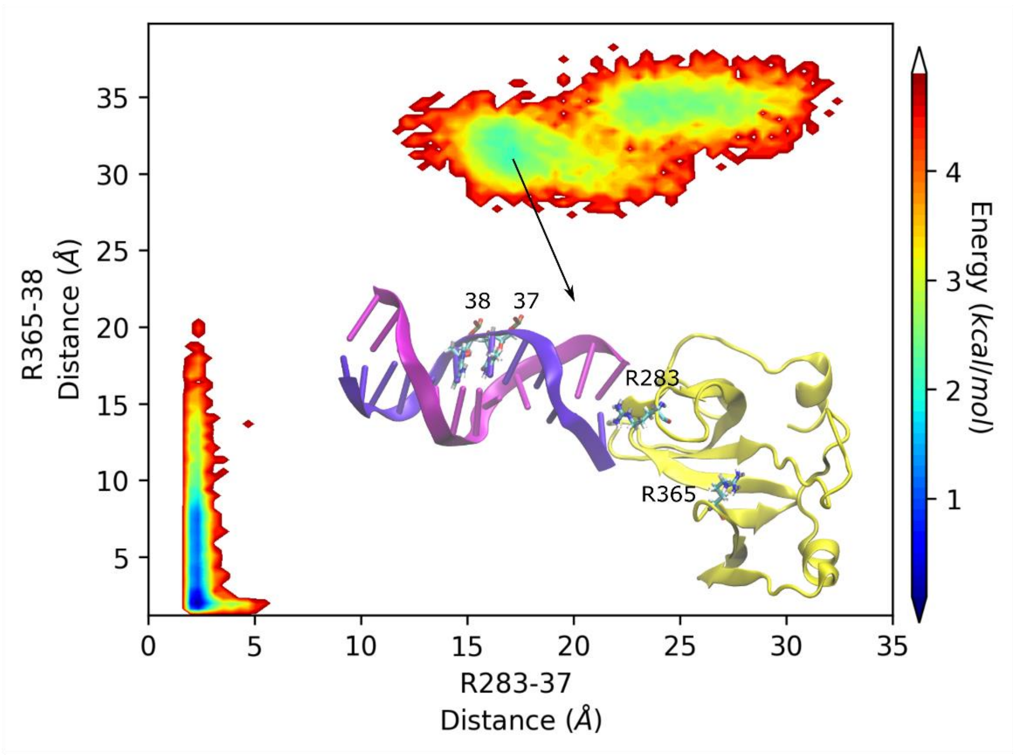


**Figure S4.** Free energy landscape for the distances between R283 of protein and residue 37 of DNAB (x-axis) and R365 of protein and residue 38 of DNAB (y-axis) with the AMBER ff14sb-bsc1 force field. The snapshot shows the conformation at the minimum energy of one replicate simulation in which protein and DNA lost most of their contacts.

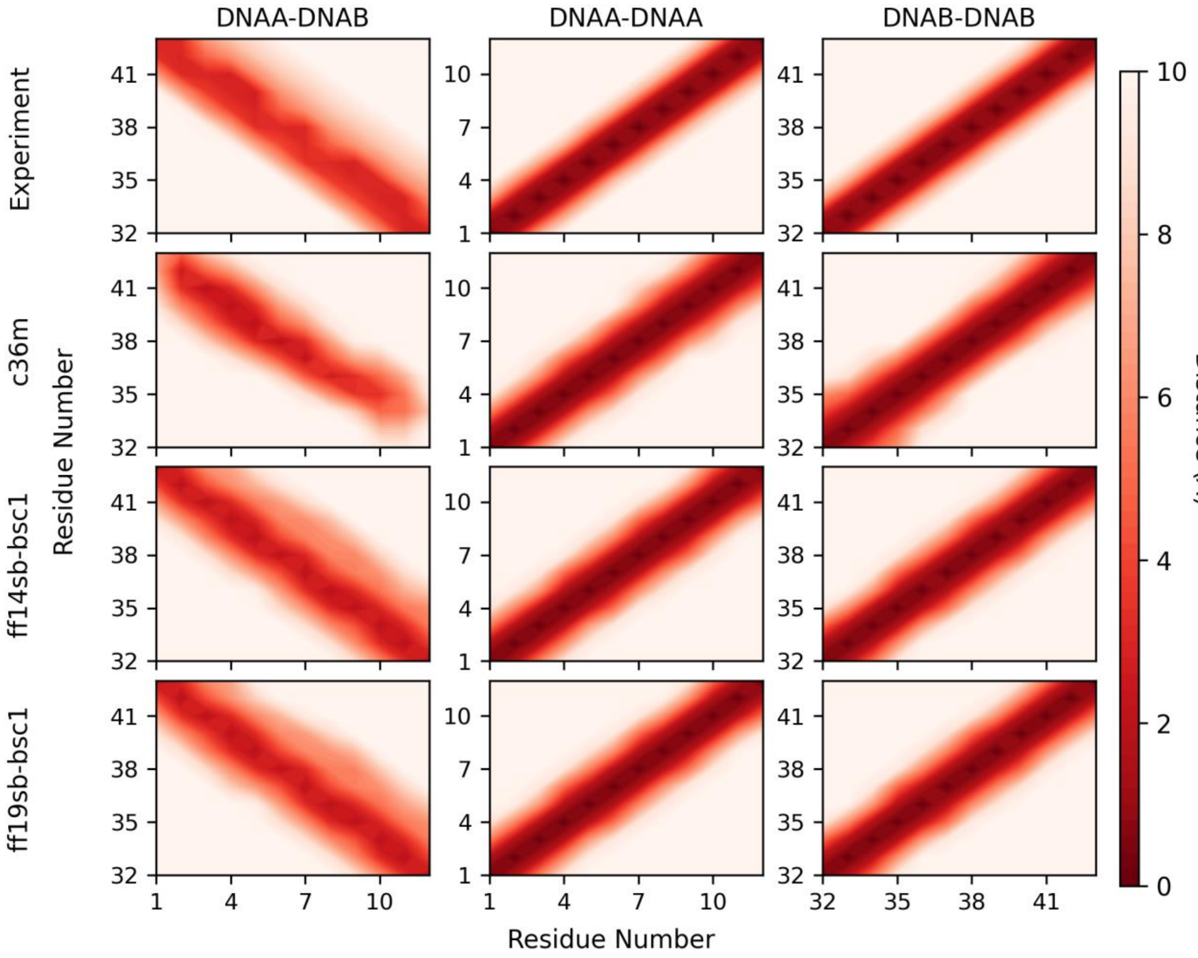


**Figure S5.** Distance maps analysis for the simulations of the KOW1-DNA system with CHARMM c36m, AMBER ff14sb and AMBER ff19sb force fields. Distance maps between DNA chains (DNAA-DNAB, DNAA-DNAA, and DNAB-DNAB) were calculated from the distances for the experimental structure (PDB ID:5oik) and average distances over the trajectories.
